# Considerations for metabarcoding-based port biological baseline surveys aimed at marine non-indigenous species monitoring and risk-assessments

**DOI:** 10.1101/689307

**Authors:** Anaïs Rey, Oihane C. Basurko, Naiara Rodriguez-Ezpeleta

## Abstract

1. Monitoring introduction and spread of non-indigenous species via maritime transport and performing risk assessments require port biological baseline surveys. Yet, the comprehensiveness of these surveys is often compromised by the large number of habitats present in a port, the seasonal variability and the time-consuming morphological approach used for taxonomic identification. Metabarcoding represents a promising alternative for rapid comprehensive port biological baseline surveys, but its application in this context requires further assessments.
2. We applied metabarcoding (based on barcodes of the Cytochrome c oxidase subunit I and of the 18S ribosomal RNA gene) to 192 port samples collected i) from diverse habitats (water column – including environmental DNA and zooplankton, sediment and fouling structures), ii) at different sites (from inner to outer estuary), and iii) during the four seasons of the year.
3. By comparing the biodiversity metrics derived from each sample group, we show that each sampling method resulted in a distinct community profile and that environmental DNA alone cannot substitute for organismal sampling, and that, although sampling at different seasons and locations resulted in higher observed biodiversity, operational results can be obtained by sampling selected locations and seasons.
4. By assessing the taxonomic composition of the samples, we show that metabarcoding data allowed the detection of previously recorded non-indigenous species as well as to reveal presence of new ones, even if in low abundance.
5. *Synthesis and application*. Our comprehensive assessment of metabarcoding for port biological baseline surveys sets the basics for cost-effective, standardized and comprehensive monitoring of non-indigenous species and for performing risk assessments in ports. This development will contribute to the implementation of the recently entered into force International Convention for the Control and Management of Ships’ Ballast Water and Sediments.

## Introduction

Globalization has led to an increased maritime transportation, with more and larger ships than ever before transferring species into ports via ballast water and hull fouling (Seebens, Gastner, & Blasius, 2013). Among the thousands species arriving daily (Carlton, 1999), some are Non-Indigenous Species (NIS) and can become invasive, disrupting native populations (Bax, Williamson, Aguero, Gonzalez, & Geeves, 2003). Thus, being important gateways for introduction of NIS, ports need to be monitored to provide information required by legal frameworks aiming at controlling biological invasions (Lehtiniemi et al., 2015).

One of these legal frameworks is the International Convention for the Control and Management of Ships’ Ballast Water and Sediments (IMO, 2004), which requires that ships treat their ballast water before its release to port, unless they show that the risk of transferring NIS between the donor and recipient ports is limited (David, Gollasch, & Pavliha, 2013). Such risk assessment requires cataloguing biodiversity through Port Biological Baseline Surveys (PBBS), which requires collecting samples using various methods (Kraus et al., 2018) and at different spatial and temporal scales (Lehtiniemi et al., 2015) given the diverse range of habitats (such as soft sediment, water column, or exposed and sheltered artificial structures) allowing presence of distinct organismal groups (such as benthic macrofauna, fouling organisms or planktonic organisms).

Global initiatives have been initiated to standardize sampling for PBBS (HELCOM/OSPAR, 2013; Awad, Haag, Anil, & Abdulla, 2014; Kraus et al., 2018) and are based on morphological taxonomic identification of the species found. Yet, this method lacks accuracy for identification of larvae and eggs, developmental stages at which many NIS are transported (Gittenberger, Rensing, Niemantsverdriet, Schrieken, & Stegenga, 2014) and relies on taxonomists who are often specialized on the local biota, but who have limited knowledge of alien taxa (Pyšek et al., 2013) and/or on specific taxonomic groups coexisting in a port (Bishop & Hutchings, 2011). Moreover morphological identification is time-consuming (Mandelik, Roll, & Fleischer, 2010), especially when considering the several tens of samples required to characterize a port. These limitations translate into PBBS being completed several years after sample collection (Bott, 2015), reducing the effectiveness of prevention of NIS introduction and spread control strategies.

DNA metabarcoding (Taberlet, Coissac, Pompanon, Brochmann, & Willerslev, 2012), the simultaneous identification of taxa present in a complex environmental sample based on a conserved DNA fragment, is revolutionizing traditional biodiversity monitoring (Creer et al., 2016; Elbrecht, Vamos, Meissner, Aroviita, & Leese, 2017), and could represent the cost-effective and possible to standardize alternative required for PBBS (Lehtiniemi et al., 2015). Metabarcoding can be applied to DNA extracted from bulk organismal samples (community DNA) or to environmental DNA (eDNA), which is extraorganismal DNA released in the environment in form of cells, feces, skin, saliva, mucus, etc. (Shaw, Weyrich, & Cooper, 2016). The later has received particular attention lately for its potential to cost-effectively and noninvasively survey species richness from many ecosystems (Deiner, Fronhofer, Mächler, Walser, & Altermatt, 2016), and for being able to detect spatially discrete communities (Jeunen, Knapp, Spencer, Lamare, et al., 2019).

Metabarcoding has been extensively explored and validated as an alternative tool to morphological identification (e.g. Aylagas et al. (2016; 2018), and described as a promising tool for NIS monitoring (Comtet, Sandionigi, Viard, & Casiraghi, 2015). Recent studies have highlighted the potential of metabarcoding applied to eDNA to perform port biodiversity surveys (Borrell, Miralles, Huu, Mohammed-Geba, & Garcia-Vazquez, 2017; Grey et al., 2018; Lacoursière-Roussel et al., 2018). Yet, very few studies have performed port surveys based on additional sampling substrates (Brown, Chain, Zhan, MacIsaac, & Cristescu, 2016; Zaiko et al., 2016) and only one has compared biodiversity assessments obtained by the different substrates (Koziol et al., 2019). Additionally, few studies have considered the spatial and temporal variability of ports (e.g. eDNA in arctic ports Lacoursière-Roussel et al. (2018) and settlement plates in austral temperate port Zaiko et al. (2016)) but none has fully evaluated the impact of spatial and temporal variability on all the different communities of a port, which is critical to ensure maximum biodiversity recovery in PBBS.

Here, we have applied metabarcoding to 192 samples collected from the port of Bilbao (Northern Spain) to evaluate the effect of i) using various sampling methods to capture biodiversity found in different substrates, ii) sampling at different sites from outer to inner estuary, iii) sampling during different seasons of the year, and iv) using alternative genetic markers. Our results provide relevant information for performing non-indigenous species monitoring and risk assessments in ports in response to the International Convention for the Control and Management of Ships’ Ballast Water and Sediments.

## Materials and methods

### Sampling

Zooplankton, sediment, fouling and water samples were collected from the port of Bilbao in autumn 2016, winter 2017, spring 2017, and late summer 2017 at four sites (Fig 1 and Appendix 1). Sites 1, 2 and 3 represent the outer parts of the port with busy berths and are characterized by deep waters (10-30 m) and salinity (>30 PSU); site 4 represent s the inner, less busy part of the port and is characterized by low water depth (6-9 m) and lower salinity (>20 PSU). Zooplankton, sediment and fouling sampling was based on the protocol designed by Helcom/Ospar (HELCOM/OSPAR, 2013). Zooplankton samples were collected in vertical tows using Pairovet nets (mesh sizes of 60 and 280 μm) at three points per site, mixing the collected material to have one sample per mesh size per site. The collected zooplankton was grinded with a mortar until no integer organism could be appreciated and was stored in 96% ethanol at −20 °C. Sediment samples (3 samples per site) were collected by sieving material collected from a Van Veen grab (0.07–0.1 m^2^) with a 1 mm mesh size sieve. Retained material was processed following Aylagas et al. (2016) based on decantation and homogenization phases, and retrieved benthic macroinvertebrates were stored in 96% ethanol at −20 °C. Homogenization was done by blending the decanted organic material in a PHILIPS hr2095 700 W 2 L glass jar with 96% ethanol until no fragments of animals and others organic materials could be observed. During decantation and homogenization steps, all laboratory equipment used were cleaned between samples by soaking in 10% bleach for 15 min and thoroughly rinsing with deionized water. Fouling samples were collected by placing 15 × 15 cm polyvinyl chloride plates at 1, 3 and 7 m depth in a suspended array in each site. Plates were deployed in winter and spring and recovered in spring and late summer respectively (see Table S1). At recovery, were placed in individual sterile plastic bags, soaked in 96% ethanol and thoroughly scrapped both sides with a scalpel to retrieve fouling organisms attached to it. The detached organisms were then homogenized with a blender (Conair™ Waring™ Mini-sample Containers for Blender) in 96% ethanol and stored in 96% ethanol at −20 °C. Cleaning procedure was similar as the one performed for sediment samples. At each sampling site, 1 L surface water sample was collected from each sampling point with a bottle. Samples from the same site and from the same sampling method were combined into a single sample by site. In sites 1 and 3, three additional samples were taken at one meter above the bottom using a Niskin bottle and combined. Each 3 L combined water sample was filtered using a 0.45 μm Sterivex filter unit (Merck Chemicals & Life Science), which was stored at −80 °C until further use. Additionally, 3 L surface water sample from three sampling sites from the ports of Vigo and A Coruña, located in Galicia, Northwestern Spain following the same sampling procedure than for Bilbao by combining 1 L water from three sampling point per sampling site, were collected in March 2017 and filtered as described above.

**Figure 1.**
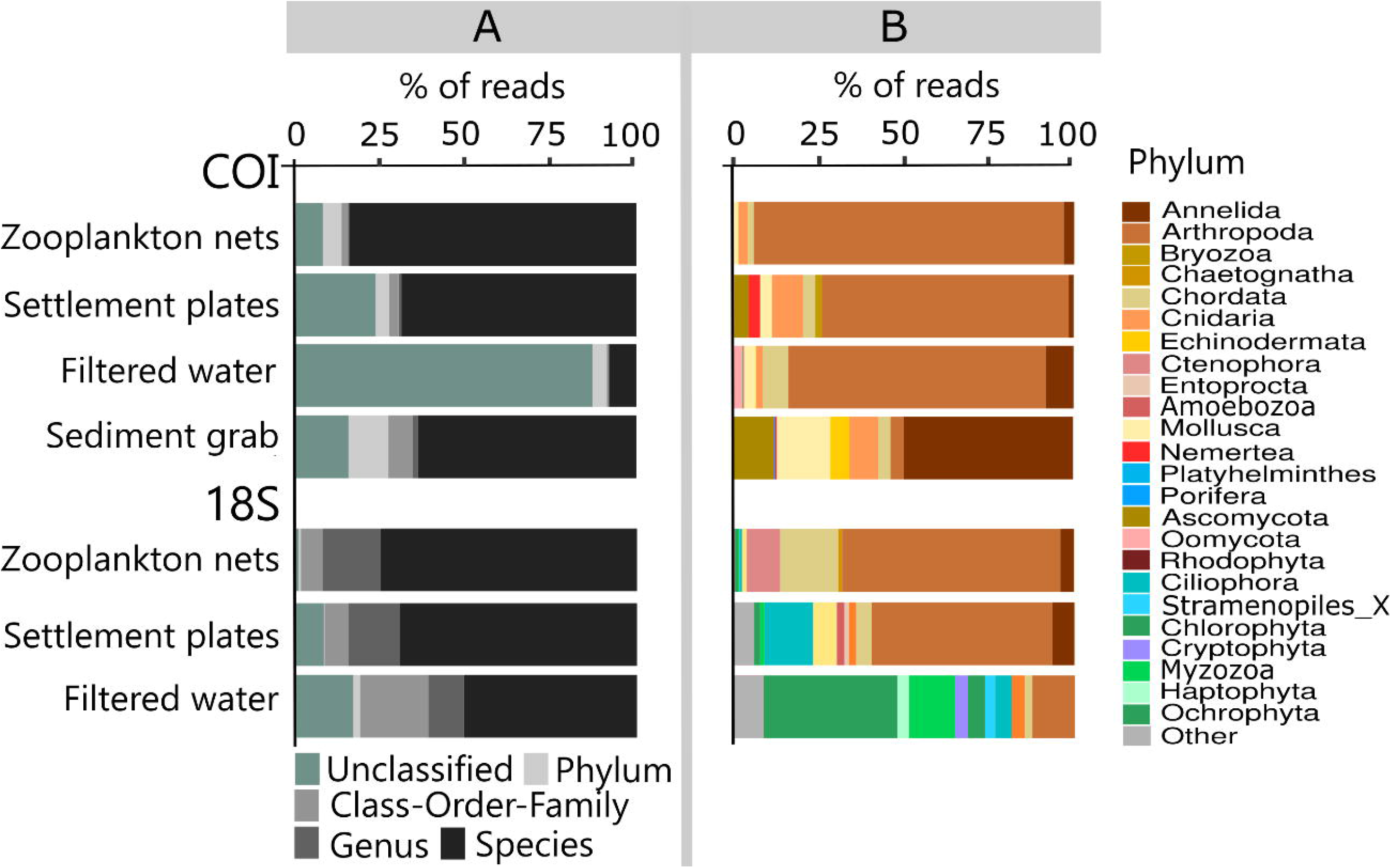
Sampling protocol representing sampling location (Port of Bilbao), sites (1, 2, 3 and 4) and points per site (dots); and illustrating the sampling methods used and the targeted biological communities.

### DNA extraction, library preparation and sequencing

Total genomic DNA was isolated from zooplankton samples with the DNeasy blood & tissue kit (QIAGEN), benthic macroinvertebrate and fouling organism samples with, respectively, PowerMax and PowerSoil DNA Isolation Kits (MOBIO) replacing the initial bead-beating step by an overnight incubation at 56°C with proteinase K (0.4 mg/ml). Filter samples were extracted with the DNeasy blood & tissue kit (QIAGEN) following the “SX filters without preservation buffer” developed by Spens et al. (2017). Negative controls (preservation buffer) were included in each batch of DNA extraction and followed the same procedure as all other samples. DNA concentration was measured with the Quant-iT dsDNA HS assay kit (Thermo Scientific) using a Qubit 2.0 Fluorometer (Life Technologies), purity was inferred from 260/280 and 260/230 absorbance ratios with the ND-1000 Nanodrop (Thermo Scientific), and integrity was assessed by electrophoresis in 0.7% agarose. Two primer pairs widely used to assess eukaryotic biodiversity (Hebert, Ratnasingham, & de Waard, 2003; Amaral-Zettler, McCliment, Ducklow, & Huse, 2009) were used: mlCOIintF/dgHCO2198 (COI primers), targeting a 313 bp fragment of the Cytochrome c oxidase subunit I (COI) gene (Meyer, 2003; Leray et al., 2013) and 1389F/1510R (18S primers), targeting a variable length fragment (87 to 186 bp) of the V9 hypervariable region of the 18S rRNA gene (Amaral-Zettler et al., 2009). DNA extracted from zooplankton, fouling organism and filter samples were amplified with both primer pairs, whereas DNA from sediment was only amplified with the COI primers. PCR amplifications were performed in two rounds. For the first PCR, 1 μl of genomic DNA (5 ng/μl) was added to a mix consisting in 5 μl of 2X Phusion Master Mix (Thermo Scientific), 0.2 μl of each primer (0.2 μM) and 2.6 μl of MilliQ water. For the 18S primers, PCR conditions consisted on an initial 3 min denaturation step at 98 °C; followed by 25 cycles of 10 s at 98 °C, 30 s at 57 °C and 30 s at 72 °C and finally 10 min at 72 °C. For the COI primers, PCR conditions consisted of an initial 3 min denaturation step at 98 °C; followed by 35 cycles of 10 s at 98 °C, 30 s at 46 °C and 45 s at 72 °C and finally 5 min at 72 °C. Negative controls of PCR (no template) were included within each set of PCRs. For each DNA extract, three PCR amplifications were performed and pooled. Once purified using AMPure XP beads (Beckman Coulter), the mixed PCR products were used as template for the generation of dual-indexed amplicons in a second PCR round following the “16S Metagenomic Sequence Library Preparation” protocol (Illumina) using the Nextera XT Index Kit (Illumina). Multiplexed PCR products were purified again using the AMPure XP beads, quantified using Quant-iT dsDNA HS assay kit and a Qubit 2.0 Fluorometer (Life Technologies), normalized to equal concentration and sequenced using the 2 × 300 paired-end MiSeq (Illumina). Reads were demultiplexed based on their barcode sequences.

### Raw read preprocessing, clustering and taxonomic assignment

After quality checking of demultiplexed paired-end reads with FASTQC (Andrews, 2010), forward and reverse primers were removed using Cutadapt (Martin, 2011) with the anchored 5’ adapter and for “paired end reads” options and with the linked adapter option for COI and 18S respectively. Forward and reverse reads were merged using PEAR (Zhang, Kobert, Flouri, & Stamatakis, 2014) with a minimum sequence overlap of 217 bp and maximum amplicon length of 313 bp and of 80 and 190 bp for COI and 18S barcodes respectively. Merged reads with average Phred quality score lower than 20 were removed with Trimmomatic (Bolger, Lohse, & Usadel, 2014). Using Mothur (Schloss et al., 2009), sequences without ambiguous bases were aligned to BOLD (https://www.boldsystems.org) or SILVA (https://www.arb-silva.de/documentation/release-132/) for COI and 18S respectively, and only those covering the barcode region were kept. Chimeras, detected using *de novo* mode of UCHIME (Edgar, Haas, Clemente, Quince, & Knight, 2011), were removed, and remaining reads were clustered into OTUs using Swarm 2.2.1 with the step-by-step aggregation clustering algorithm implemented with default settings (Mahé, Rognes, Quince, de Vargas, & Dunthorn, 2014). SWARM algorithm does not rely on a fixed threshold for delimiting OTUs which is pertinent when performing PBBS where highly diverse biodiversity can be found. “Singleton” OTUs, composed by a single read, were removed. No rarefaction was performed to avoid decreasing sensitivity by choosing an arbitrary minimum library size (McMurdie and Holmes, 2014). The remaining OTUs were taxonomically assigned using the Naïve Bayes Classifier (Wang, Garrity, Tiedje, & Cole, 2007) using BOLD (accessed in May 2018) or PR2 (release 4.10.0) databases for COI and 18S respectively.

### Community analyses

Apart from the complete dataset, we created two subsets: a taxa targeted through PBBS dataset, including only reads classified to the class level and excluding those matching to non-targeted groups for PBBS such as Mammalia, Aves, Insecta, Collembola, Arachnida and all classes of Fungi; and a NICS dataset, including only reads matching either the 68 Non-Indigenous and Cryptogenic Species (NICS) previously detected in the port of Bilbao (Martínez & Adarraga, 2006; Adarraga & Martínez, 2011, 2012; Butrón, Orive, & Madariaga, 2011; Zorita et al., 2013; Tajadura, Bustamante, & Salinas, 2016) or the 1083 species present in the AquaNIS database (AquaNIS. Editorial board., 2015). Most analyses were conducted using RStudio (Team, 2015) with *vegan* (Oksanen et al., 2013), *adespatial* (Dray et al., 2018) and *indicspecies* (De Cáceres & Legendre, 2009) libraries. Indicator species analyses (Dufrene & Legendre, 1997) were performed on 1) the taxa targeted through PBBS dataset to identify indicator taxa of each sampling method and 2) the NICS dataset to identify non-indigenous indicator taxa of each sampling site. These analyses were based on the IndVal index calculated as the product of the degree of specificity (measuring the uniqueness to a sampling method or site) and the degree of fidelity (measuring the frequency of occurrence within a sampling method or site) of an OTU to a given sampling condition. Statistical significance of associations was assessed by performing 10,000 permutations. The effects of season and locality on taxa targeted through PBBS communities were tested for significance using a permutational multivariate analysis of variance (PERMANOVA) after checking for multivariate homogeneity of group dispersions (betadisper). PERMANOVA and betadisper were performed on Euclidean distances on Hellinger-transformed OTU abundances (Hellinger distances), which are appropriate for community ordination and clustering (Legendre & Gallagher, 2001) and on Jaccard dissimilarities based on OTU presence/absence. The contribution of replacement (changes in OTU identity) and nestedness (richness differences where one sample is a subset of a richer sample) to beta diversity of taxa targeted through PBBS between seasons and between sites was computed using the relativized nestedness index of Podani and Schmera (2011) based on Jaccard dissimilarities matrix. For each sampling method, replacement and nestedness were calculated between pairwise comparisons of 1) samples belonging to the same site but sampled at different seasons (season variation of beta diversity), and 2) samples belonging to the same season but sampled at different sites (spatial variation of beta diversity). The mean proportion of nestedness and replacement contribution to beta-diversity between sites and between seasons was then calculated. For all OTUs assigned to a Non-Indigenous and Cryptogenic Species, we blasted their representative sequences against BOLD and PR2 databases respectively for COI and 18S barcodes.

## Results

### Communities retrieved by each sampling method and genetic marker

A total of 5,718,639 and 7,055,675 COI and 18S barcode reads were kept for analysis (Table S1). Negative controls showed little contamination (with only 164, 66 and 213 of COI and 2031 and 163 18S quality filtered reads). The reads corresponding to the 192 samples collected at the port of Bilbao resulted in 40,318 and 20,473 OTUs for COI and 18S respectively. For all sampling methods, OTU accumulation curves of OTUs against reads approached saturation, suggesting that adding more sequencing effort would provide limited increase in diversity (Figs 2A and Appendix 2). The majority of the OTUs, 89% for COI and 73% for 18S, were unique to one sampling method (Fig. 2B), and, for both barcodes, but particularly for 18S, filtered water (aimed at retrieving eDNA from macroorganisms and microbial eukaryotes) resulted in higher OTU richness and unique OTUs than other sampling methods (Figs 2A, B and Appendix 2). Different sampling methods retrieved distinct biological communities as observed with the PCA (Fig. 2C) and confirmed with PERMANOVA analyses (COI: R^2^=0.28, p-value= 9.99e-05; 18S: R^2^=0.30, p-value=9.99e-05).

**Figure 2.**
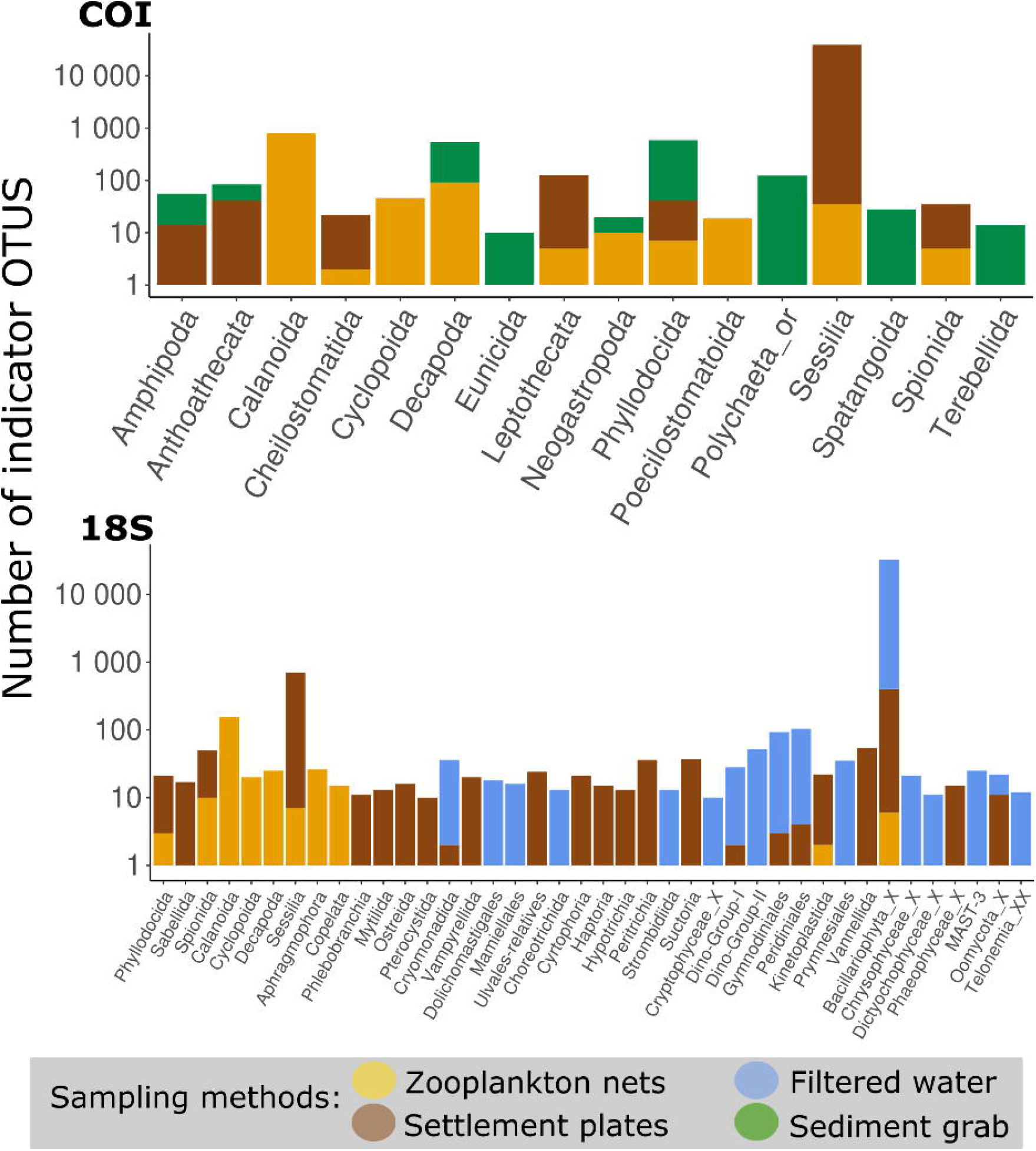
Overall description of community detected per barcode. A: OTU accumulation curves per sampling method. B: Venn diagrams of the number and percentage of OTUs shared between sampling methods. C: Principal Component Analyses of the Hellinger-transformed abundances of OTUs. Sample scores are displayed in scaling 1 with ellipses representing the 95% confidence dispersion of each sampling method.

The percentage of reads assigned to species or higher taxonomic levels differed between sampling methods and barcodes. The majority of the reads were assigned to species level for zooplankton nets (COI: 84% and 18S: 75%), settlement plates (COI: 69% and 18S: 69%) and sediment grabs (COI: 64%); for filtered water, although 51% of reads were assigned to species level with 18S, only 8% could be assigned to a species with COI (Fig. 3A). Filtered water analyzed with COI had also the largest proportion of reads that could not be assigned to phylum (87%). This was due to the non-specific amplification of prokaryotic and non-target eukaryotic DNAs (Appendix 3), which is known for COI (Collins et al., 2019).

**Figure 3.**
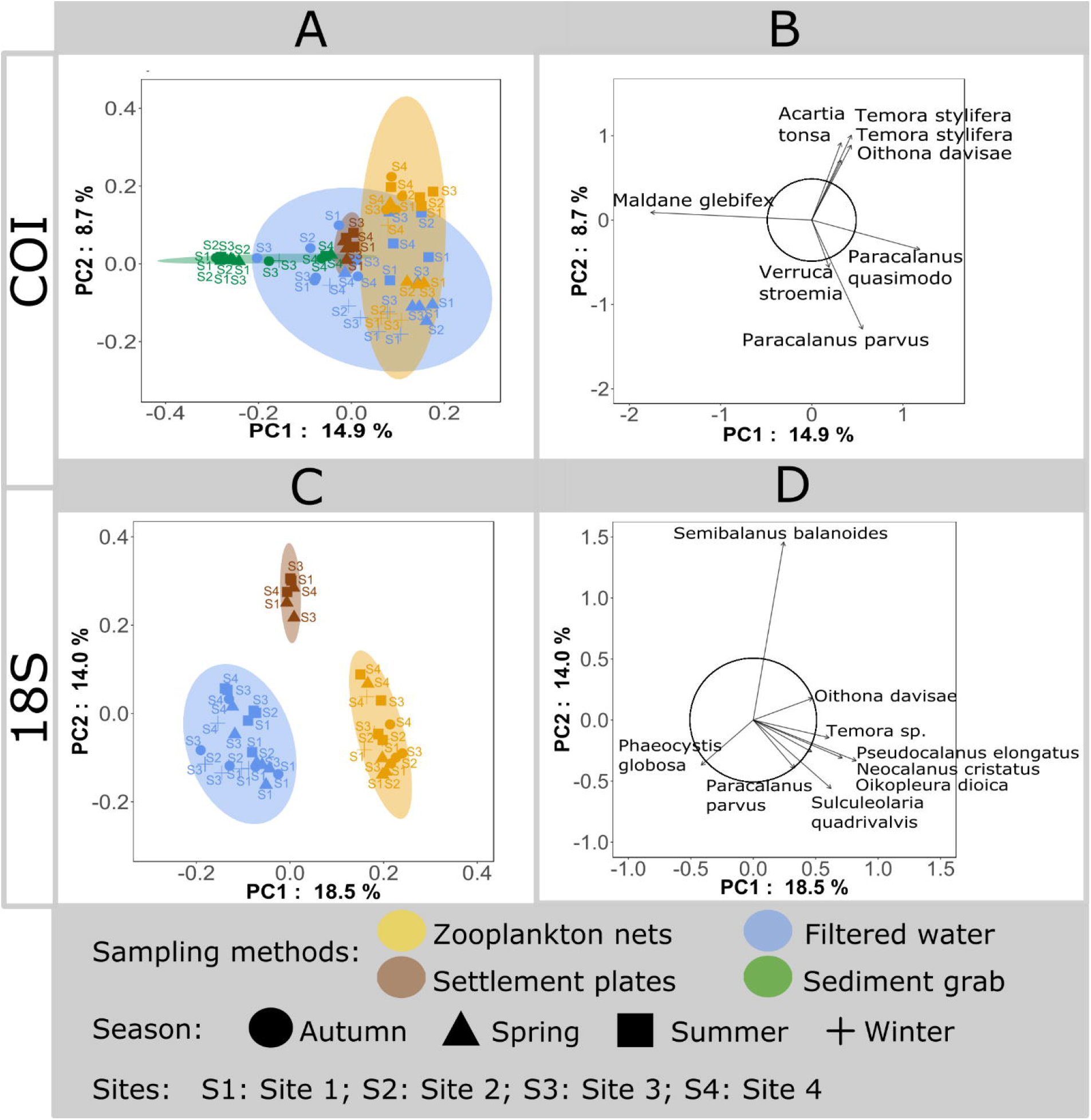
Taxonomic assignment per barcode and per sampling method. A: Percentage of reads assigned to each taxonomic level. B: Relative abundance of reads classified to at least Phylum level for the 10 most abundant phyla.

Both barcodes detected a wide range of eukaryotic groups (Fig. 3B). With COI, the greatest majority of reads belonged to metazoans, with Arthropoda dominating all sampling methods except sediment grabs, dominated by Annelida. With 18S, a more diverse spectrum of taxa was retrieved, including phytoplankton and macroalgae; as expected, the number of reads assigned to these phyla in zooplankton nets and settlement plates was very low in comparison to filtered water.

### Distribution of taxa targeted through port biological baseline survey

A total of 16,828 and 9,091 OTUs were kept as taxa targeted through PBBS for COI and 18S respectively. From those, indicator analysis identified respectively 2,600 (15%) and 1,700 (19%) OTUs significantly associated to one of the sampling methods. Settlement plates were associated with 1,268 (COI) and 808 (18S) OTUs, zooplankton nets with 1,001 (COI) and 315 (18S) OTUs, and sediment grabs with 238 (COI) OTUs. Only 12 OTUs (9 of which were metazoans) were indicators of filtered water with COI while 580 (3 of which were metazoans), were associated to this sampling method with 18S. We observed expected strong associations of some taxa to one particular sampling method: fouling bivalves (Ostreoida, Mytilida) with settlement plates, copepods (Calanoida, Cyclopoida) with zooplankton nets and sea urchins (Spatangoida) with sediment grabs (Fig. 4). Yet, less obvious associations were also observed such as barnacles (Sessilia) and two polychaeta orders (Spionida, Phyllodocida) with zooplankton nets or dinoflagellates (Peridiniales and Gymnodiniales) with settlement plates, illustrating the advantages of a complementary sampling approach to recover the diversity of these taxonomic groups.

**Figure 4.**
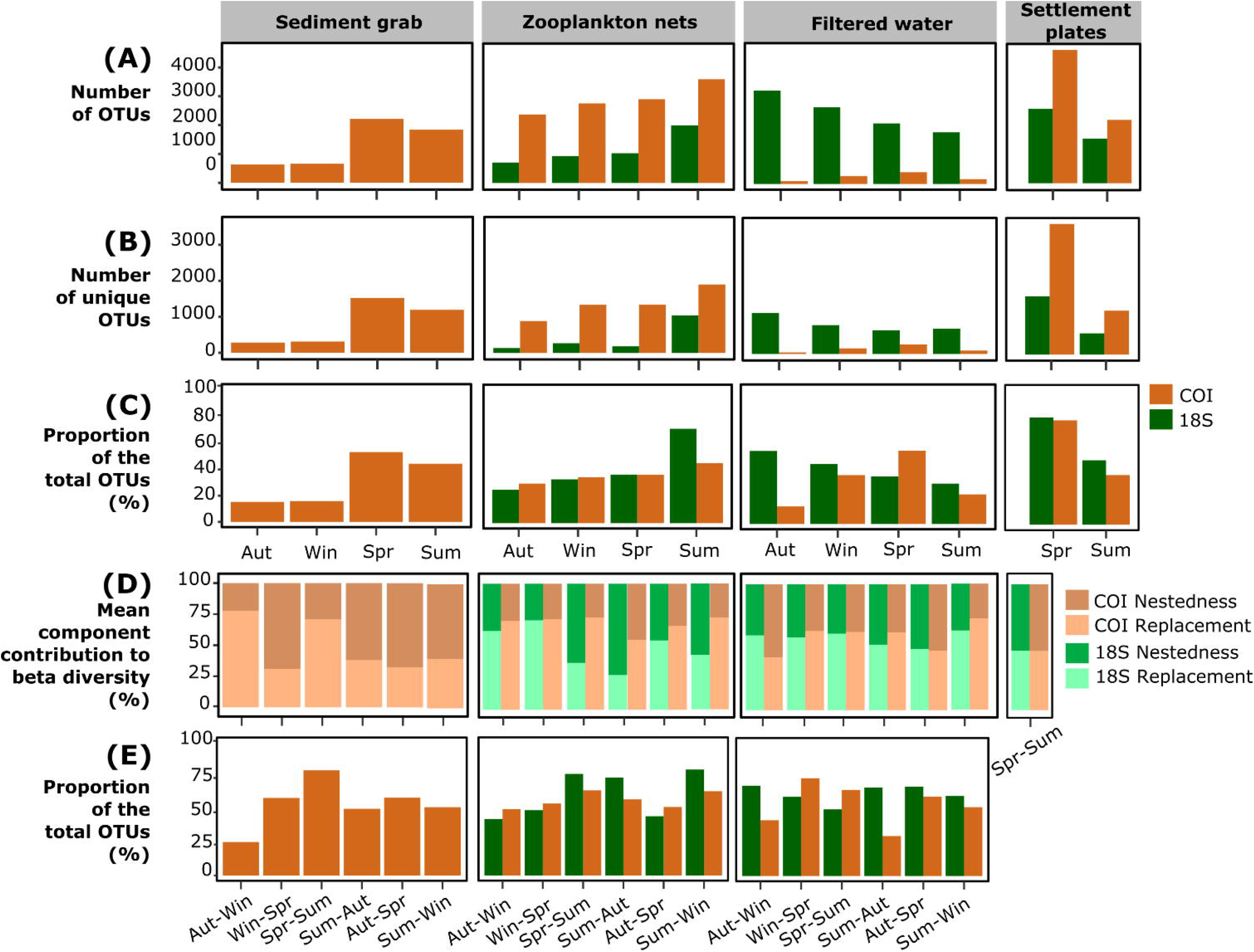
Distribution of indicator OTUs associated to each sampling method at the order level (in log scale) for COI and 18S. Only orders with at least 10 indicator OTUs are shown.

The need for complementarity of sampling methods was further confirmed by PCA on PBBS target taxa (Fig. 5). For both barcodes, expected taxonomic differences were found between the sampling methods: sediment grabs were characterized by the polychaeta *Maldane glebifex* (Fig. 5B), zooplankton nets were distinguished by several copepods (Fig. 5B, D) and settlement plates, by the encrusting *Semibalanus balanoides* (Fig. 5D). Filtered water communities were characterized differently according to each barcode. For COI, filtered water samples were not differentiated as a distinct group and were in general close to those retrieved with zooplankton samples collected at the same season (Fig. 5A). In contrast, for 18S, filtered water was different due to the presence of the phytoplankton species *Phaeocystis globosa* (Fig. 5D); yet, when targeting only metazoan taxa, the patterns observed with COI were similarly to the ones retrieved with 18S (Appendix 4). For filtered samples, the proportion of metazoan OTUs detected was largely different compared to the other methods (Appendix 5). For instance, Decapoda orders had a low diversity with filtered water (COI: 0,7%; 18S: 7%), whereas more diversity could be detected for Calanoida (COI: 8%; 18S: 35%), Sabellida (COI: 15%; 18S: 28%) and Leptothecata (COI: 13%; 18S: 58%). In general, filtered water recovered the smallest metazoan diversity.

**Figure 5.**
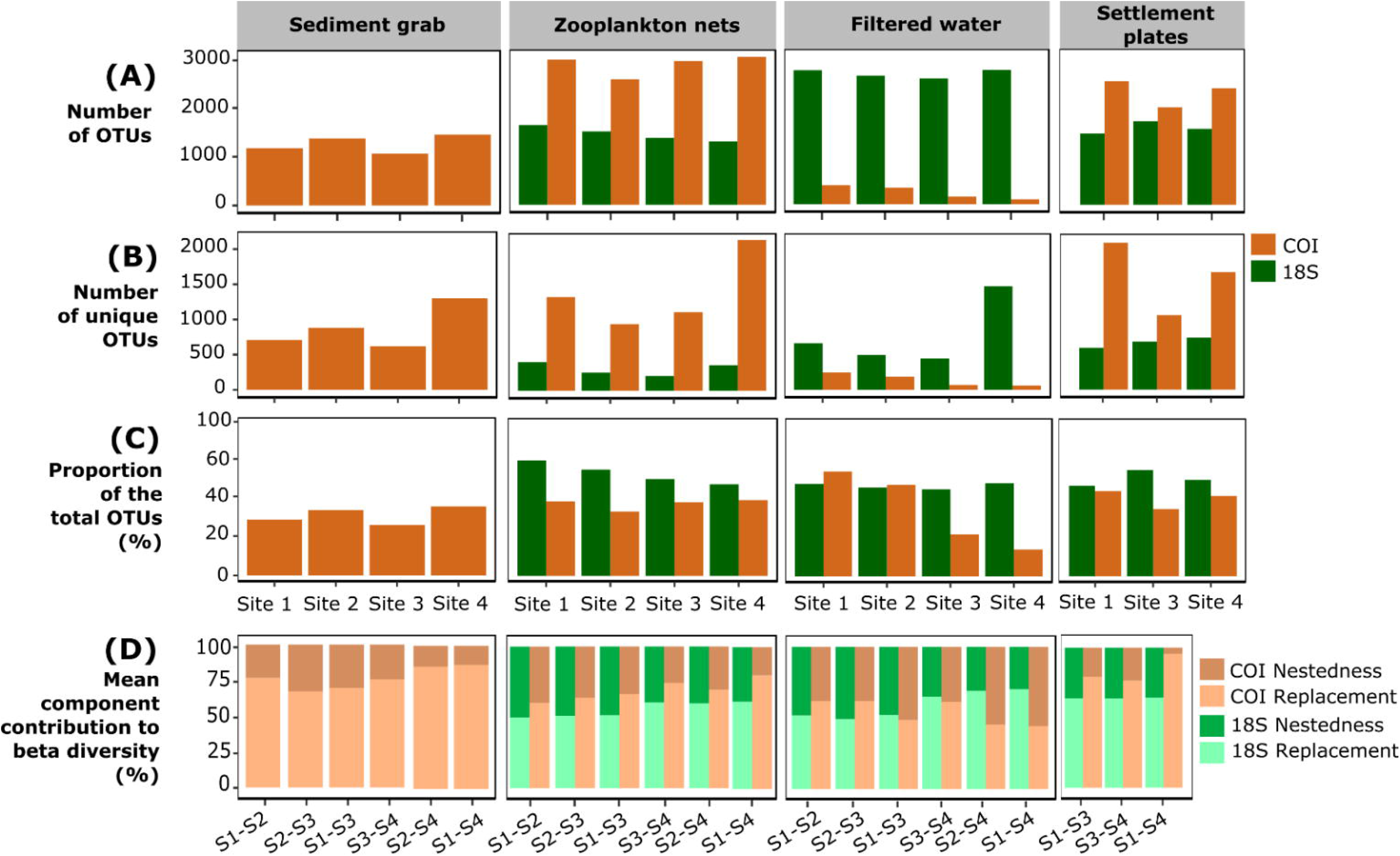
Principal Component Analyses of Hellinger-transformed abundances of OTUs included in the *PBBS targeted taxa dataset* for COI (A-B) and 18S (C-D). A & C: Samples scores in scaling 1 with ellipses representing the 95% confidence dispersion of each sampling method. B & D: OTU scores in scaling 1 with the circle of equilibrium contribution. Only OTUs whose projected length in these two principal component axes exceeding the value of equilibrium contribution are represented.

### Influence of sampling seasonality and locality on detected biodiversity

Seasonal variations contributed significantly to differences in community composition of zooplankton nets and filtered water but not of sediment grab (Table S2). Total richness and unique richness varied between seasons for all sampling methods (Fig. 6A, B). Spring and late summer were generally richer than autumn and winter, excepted for filtered water communities represented with 18S. For sediment grabs and zooplankton nets with 18S, a strong proportion of taxa found in autumn and winter were a subset of taxa retrieved in late summer and spring. Indeed, the nestedness component of sediment grab assemblages represented between 60 to 70% of total compositional variations among these pairs of seasons (Fig. 6D). For zooplankton net assemblages, 63 and 72% of nestedness was observed between late summer and spring and between late summer and autumn respectively, suggesting that late summer recovered a majority of spring and autumn diversity (Fig. 6D). Thus, the combination of spring and late summer for sediment grabs retrieved 85% of the total OTU richness over the four seasons, while for zooplankton nets, in late summer alone 71% was retrieved (Fig. 6C, E). For filtered water and settlement plates, seasonal community changes were driven by both OTU replacement and nestedness, with similar relative contributions (Fig. 6D). The seasonal influence observed in intra-port samples was also observed between ports (Appendix 6). The differences in communities between Bilbao, A Coruña and Vigo (ports belonging to the same ecoregion but separated by over 500 km) during the same season was generally smaller than that between seasons in the same port, indicating that communities were driven by seasonality rather than location (Appendix 6 and Table S3). This pattern was more pronounced with 18S than with COI.

**Figure 6.**
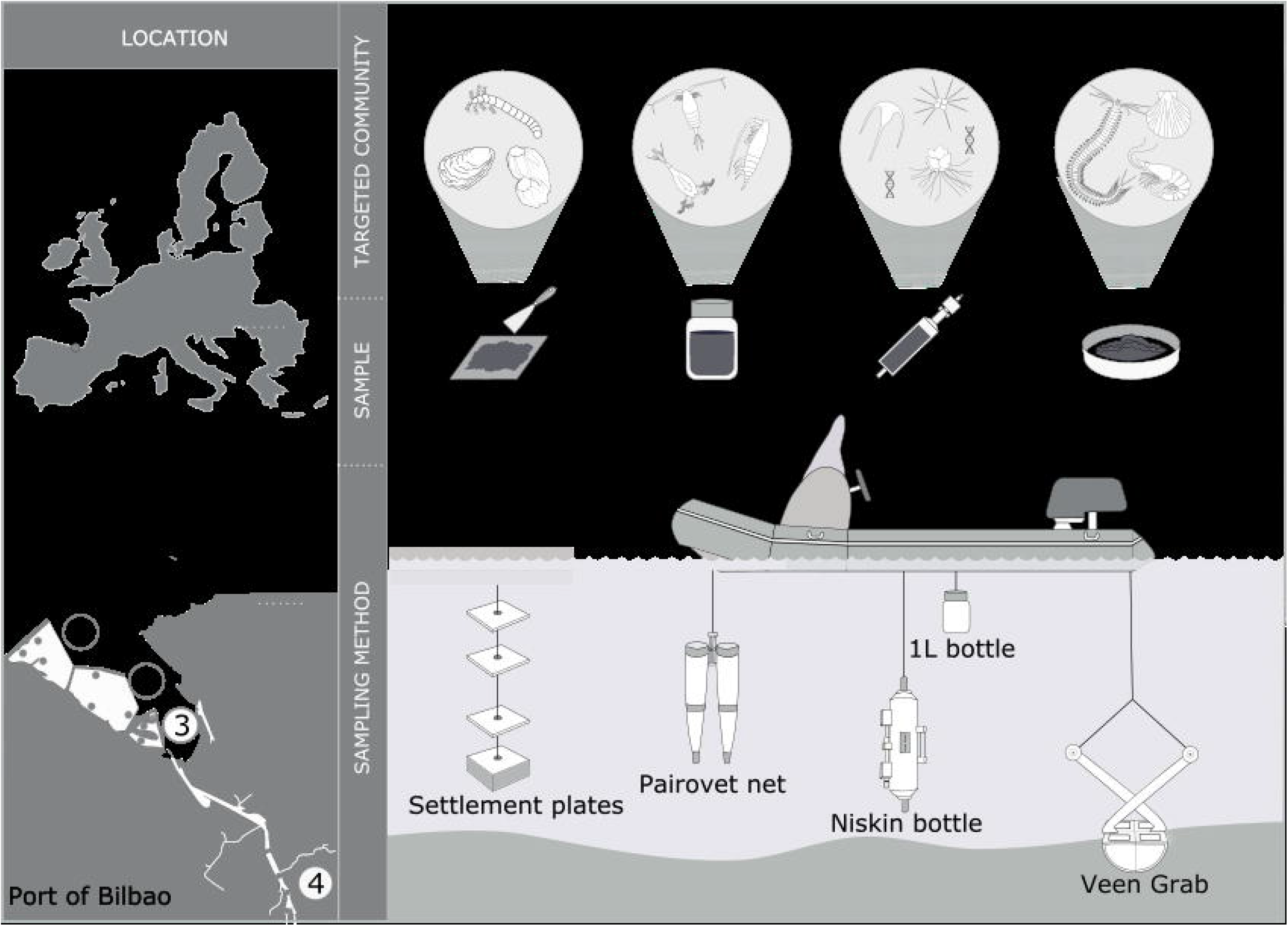
Seasonal variation of alpha and beta diversity for each sampling method with COI and 18S. A: Total OTU richness recovered at each season. B: OTU richness unique to each season. C: Proportion of the total OTU richness detected with one season. D: Decomposition of between-season beta diversity into replacement and nestedness components. E: Proportion of the total OTU richness detected with two seasons.

Locality within the Bilbao port appeared to impact on benthic assemblages (Table S2), since sites outside the estuary (sites 1 to 3) were different from site 4 inside the estuary; mainly, the polychaete *Maldane glebifex* was less abundant in site 4 (Fig. 5B). Regarding zooplankton nets, site 4 was the main driver of difference in communities as, when not considered, no significant differences between sites were observed (Table S2). Each site harbored a similar proportion of the total OTU richness found by each sampling method (Fig. 7C). This proportion did not exceed 60%. Indeed, OTU replacement contributed more to community variation among sites than nestedness, especially with sediment grab and settlement plates (Fig. 7D). OTU replacement was more important when comparing sites from outside the estuary (sites 1, 2 and S) with site 4 inside the estuary, while it contributed less to community variation among sites 1,2 and 3. This is congruent with site 4 having generally more unique OTUs than the other sites (Fig. 7B). An exception was observed for filtered water with COI, where site 4 had the lowest OTU richness and unique OTU richness, and where nestedness contributed more to the variation in community composition in comparison to sites 1 and 2 (Fig. 7A, B, D).

**Figure 7.**
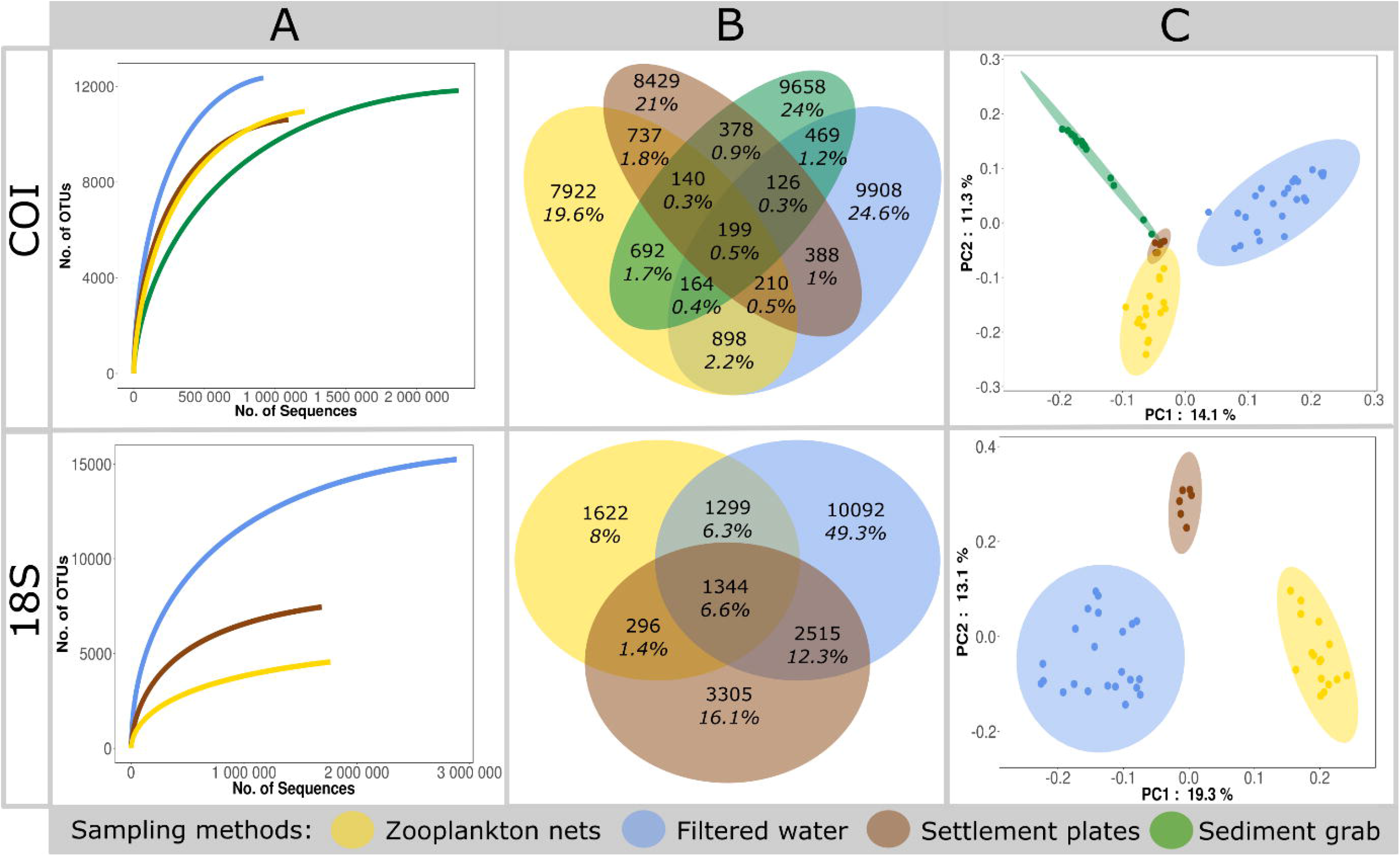
Spatial variation of alpha and beta diversity for each sampling method with COI and 18S. A: Total OTU richness recovered at each site. B: OTU richness unique to each site. C: Proportion of the total OTU richness detected with one site. D: Decomposition of between-site beta diversity into replacement and nestedness components

### Detection of Non-Indigenous and Cryptogenic Species (NICS)

Our port baseline biological survey detected 79 putative NICS, among which 29 of the 68 previously recorded in the port Bilbao were found (Tables S4, 5 and 6). Most of the other species (43) were previously detected NICS inside the port’s Large Marine Ecoregion (LME) but, for 7 of them (*Ammothea hilgendorfi*, *Bugulina fulva*, *Grandidierella japonica*, *Melita nitida*, *Neodexiospira brasiliensis*, *Pseudochattonella verruculosa*, *Tubificoides pseudogaster*), it was, to our knowledge, the first report in the LME. The indicator species analysis performed on reads corresponding to non-indigenous species revealed that site 4 was associated to the highest number of non-indigenous and cryptogenic species, which were *Oithona davisae*, *Acartia tonsa* and *Allita succinea*, found by both barcodes, *Ficopomatus enigmaticus*, found with 18S and *Amphibalanus eburneus*, *Grandidierella japonica*, *Polydora cornuta*, *Austrominius modestus*, *Monocorophium acherusicum*, *Xenostrobus securis*, found with COI. Regarding the three other sites, for COI, site 1 was associated to *Clytia hemisphaerica* and to *Balanus trigonus*, site 2, to *Clytia hemisphaerica* and *Mytilus edulis* and site 3, to *Balanus trigonus*. For 18S, only site 1 was associated to *Gymnodinium aureolum*.

## Discussion

Metabarcoding-based port baseline surveys require a combination of sampling methods and should not rely solely on eDNA

Most of the past attempts on using metabarcoding for port monitoring have relied on a single sampling method (Brown et al., 2016; Zaiko et al., 2016; Borrell et al., 2017; Grey et al., 2018; Lacoursière-Roussel et al., 2018) and only one evaluated the importance of using different sampling methods (Koziol et al., 2019). In agreement with the latter, our metabarcoding analysis shows that each sampling method recovered a distinct subset of the port community, and that, despite some taxa being expectedly associated to a given sampling method, the total diversity, even within specific taxonomic groups, was only recovered by combining different methods. Interestingly, despite the documented potential of eDNA metabarcoding to capture a large fraction of the macroorganismal diversity with limited effort (Bista et al., 2017; O’Donnell et al., 2017), including detection of non-indigenous taxa (Klymus, Marshall, & Stepien, 2017) and port surveys (Borrell et al., 2017; Grey et al., 2018; Lacoursière-Roussel et al., 2018), our analyses show that, compared to bulk sample metabarcoding, eDNA metabarcoding recovered only a subset of the metazoan diversity and did not provide additional information on targeted groups. This confirms previous findings in coral reef sites (Djurhuus et al., 2018) and freshwater streams (Macher et al., 2018) and recently in ports (Koziol et al., 2019), suggesting that despite requiring less sampling effort, COI and 18S based metabarcoding of eDNA obtained from filtered water should not be used to replace the need of obtaining bulk samples (using zooplankton nets, grabs and settlement plates) for PBBS taxa detection. Increasing evidences showed that the use of universal and degenerated COI primers is leading to non-specific amplification of prokaryotic and non-target eukaryotic DNA (Collins et al., 2019). Yet, in this context, at least two alternatives are possible to improve eDNA-based biodiversity assessments: i) increasing sequencing depth (Grey et al., 2018) or ii) using group specific markers (Jeunen, Knapp, Spencer, Taylor, et al., 2019). Yet, none of them ensures full biodiversity recovery and both significantly increase costs. Thus, the decision between increasing sampling depth and/or using multiple group specific primers for eDNA and adding multiple substrate sampling needs to be carefully considered and will be an important area of future research to optimize metabarcoding-based port monitoring.

### Metabarcoding-based port baseline surveys should include spatiotemporal sampling

While it is expected that increased temporal coverage will retrieve more taxa, the HELCOM/OSPAR protocol, a widely applied protocol developed for PBBS (http://jointbwmexemptions.org/ballast_water_RA/apex/f?p=104:13), limits sampling to late summer for sediment and fouling, and to spring and late summer for plankton in order to reduce costs. Yet, so far, no studies using metabarcoding have been performed to support this decision. Here we show that for sediment and fouling, spring sampling provides higher diversity than late summer. Although sampling in late summer could be appropriate for morphological taxonomy because of the more abundance of adult individuals, for metabarcoding, sampling in spring is preferable because during this season i) sizes of organisms are less variable and thus metabarcoding is less likely to under detect small organisms (Elbrecht, Peinert, & Leese, 2017) and ii) species diversity is at its maxima due to being a high recruitment period with abundant organisms at early life stages (Bijleveld et al., 2018). For zooplankton diversity, our results show that the HELCOM/OSPAR sampling in spring and late summer produces the highest diversity. However, the data obtained from filtered water were inconclusive. Previous studies have already shown that seasonal variations are important considerations for eDNA studies (Lacoursière-Roussel et al., 2018), but further characterizations are needed over multiple years to design an adequate protocol for maximizing biodiversity recovery with eDNA.

Concerning spatial sampling, for all sampling methods, OTU replacement generally contributed more to community variation among sites than nestedness, suggesting that spatially comprehensive sampling is crucial to recover the port’s biodiversity. Interestingly, site 4 was not only the most different from all four sites but was also the one that recorded the largest number of NIS. In coherence with our findings, it has been observed that brackish environments favors NIS settlement (Zorita et al., 2013) because these species usually support wider range of salinity (Cardeccia et al., 2018). Thus, samples for port monitoring should include those with a wide range of abiotic conditions and covering priority sampling, such as highly active ship berths, potential reservoirs of newly arrived NIS (Hewitt & Martin, 2001).

### Metabarcoding provides valuable information on Non-Indigenous and Cryptogenic Species (NICS)

Our metabarcoding port biological baseline survey detected NICS previously recorded in the port of Bilbao and NICS known to be present in the port’s marine ecoregion. Importantly, it also unveiled presence of seven NICS for which no records exist so far in this ecoregion, highlighting the potential of metabarcoding for early NICS detection. Nonetheless, not all NICS species previously recorded in Bilbao were found by our analyses. This might be due to these species not being present in the port at the time of our survey, to biases of the metabarcoding process such as differential DNA extraction, primer non-specificity, etc (Xiong, Li, & Zhan, 2016), or to database incompleteness. Primer bias and/or database incompleteness could also explain why 35 and 42% of NICS were uniquely detected with COI and 18S respectively. Yet, from the total species included in the AquaNIS database (n=1083), only 460 and 369 are included in the BOLD and PR2 databases respectively. This stresses the need of increasing reference databases and of using multiple universal primers for species detection (Grey et al., 2018). Importantly, our study confirmed that metabarcoding can detect species occurring at low abundance (Pochon, Bott, Smith, & Wood, 2013) as we found the non-indigenous amphipod *Melita nitida* with only 18 reads while it took intensive surveys in 2013, 2014 and 2016 in 3 distinct sampling regions of the Bay of Biscay to record 76 individuals (Gouillieux, Lavesque, Blanchet, & Bachelet, 2016).

## Conclusions

The Ballast Water Management Convention, entered into force in September 2017, aims at preventing the spread of non-indigenous species from one region to another (IMO, 2004). Yet, the implementation of this convention still requires technological developments for assessing compliance and granting exemptions, for which genetic methods have been suggested promising (Rey, Basurko, & Rodríguez-Ezpeleta, 2018). Based on the comparative analysis of 192 samples assessing the use of alternative sampling methods, of sampling at different seasons and at different port locations, we have demonstrated the suitability of metabarcoding for port biodiversity surveys and NIS monitoring and settled the guidelines for future studies. We show that i) combining two pairs of universal primers provides a more holistic view of the port biodiversity, ii) a combination of sampling methods is necessary to recover the different taxonomic groups, iii) environmental DNA cannot replace traditional sampling; vi) sampling should take place in spring and late summer preferably and v) spatial coverage should cover the port’s salinity gradient. Considering the cost-effectiveness of metabarcoding with respect to morphological identification, these guidelines and considerations are particularly relevant for performing the risk assessment required for granting exemptions within the International Convention for the Control and Management of Ships’ Ballast Water and Sediments.

## Acknowledgments

This manuscript is a result of the Aquainvad-ED (AQUAtic INVADers: Early Detection, Control and Management - www.aquainvad-ed.com) project funded by the European Union (Horizon 2020 research and innovation program under the Marie Sklodowska-Curie grant agreement no 642197). We thank the Port Authority of Bilbao for authorizing sampling in the port and Manuel González for making the necessary arrangements for it. We also thank Iñaki Mendibil and Elisabete Bilbao for laboratory work, Ekaitz Erauskin, Iker Urtizberea and Lander Larrañaga Juarez for sampling, Anders Lanzen for providing comments on the manuscript, and Carlota Bermejo, student of the postgraduate course in Scientific Illustration of the UPV/EHU, for producing Figure 1. This manuscript is contribution X [to be completed upon acceptance] from the Marine Research Division of AZTI.

## Authors’ contributions

AR, OB and NR-E conceived the study and designed methodology; AR and OB collected the samples; AR preformed laboratory analyses; AR and NR-E analyzed the data; AR and NR-E wrote the manuscript. All authors contributed critically to the drafts and gave final approval for publication.

## Data Accessibility

Raw demultiplexed MiSeq reads are available at NCBI SRA Bioproject PRJNA515494. Scripts used for OTU table generation and taxonomic assignment are available at https://github.com/rodriguez-ezpeleta/metabarcoding-pbbs.

## Appendix figure legends

Appendix 1: Picture of the port depicting location of each site sampled. A: Examples of the different types of structures at each site. B: Fouling communities settled on plates after 3 months. Picture frames are colored according to the site they were taken at.

Appendix 2: OTU accumulation curves per sample for COI and 18S barcodes.

Appendix 3: Number of filtered water reads assigned to each phylum for three percent identity ranges when blasting the unclassified reads (*i.e.* reads not classified below the phylum level) against NCBI. The 5 most abundant phyla per percent identity range are represented, the remaining being grouped under “other”.

Appendix 4: Principal Component Analyses of the Hellinger-transformed abundances of OTU included in the PBBS targeted metazoan taxa dataset for 18S. A: Samples scores in scaling 1 with ellipses representing the 95% confidence dispersion of each sampling method. B: OTU scores in scaling 1 with the circle of equilibrium contribution. Only OTUs whose projected length in these two principal component axes exceeding the value of equilibrium contribution are represented.

Appendix 5: Percentage of the total OTU richness detected by each sampling method for the 10 most diverse metazoan orders of the PBBS targeted taxa dataset for each barcode.

Appendix 6: Principal Component Analyses (PCA) of the Hellinger-transformed abundances of OTUs included in the PBBS targeted taxa dataset of Vigo, A Coruña and Bilbao filtered water samples. Sample scores are displayed in scaling 1. A: PCA performed with 18S. B: PCA performed with 18S targeting only metazoan. C: PCA performed with COI.

