## Supplementary figures and images for "Considerations for metabarcoding-based port biological baseline surveys aimed at marine non-indigenous species monitoring and risk-assessments"

### Appendix 1

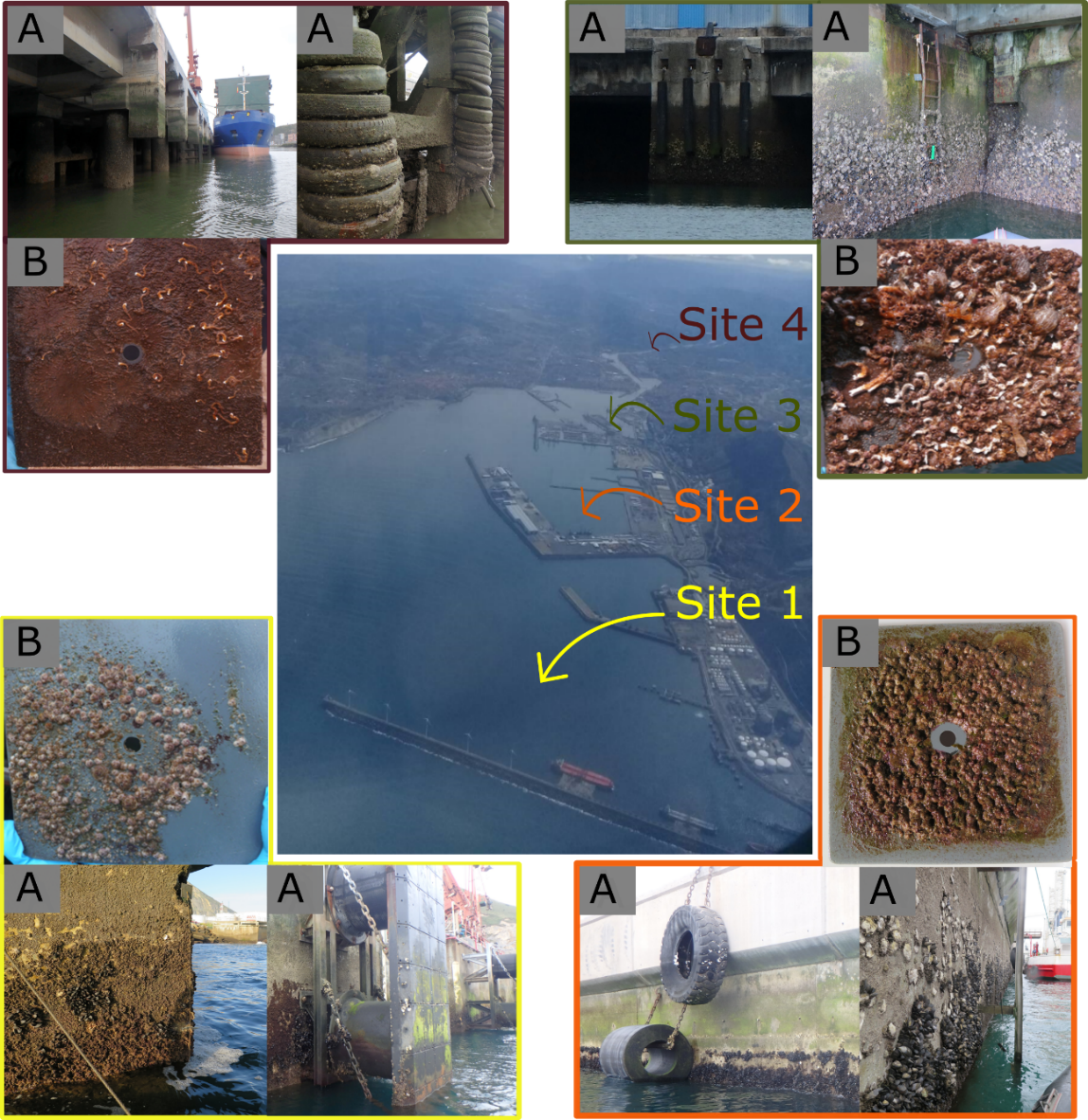

### Appendix 2

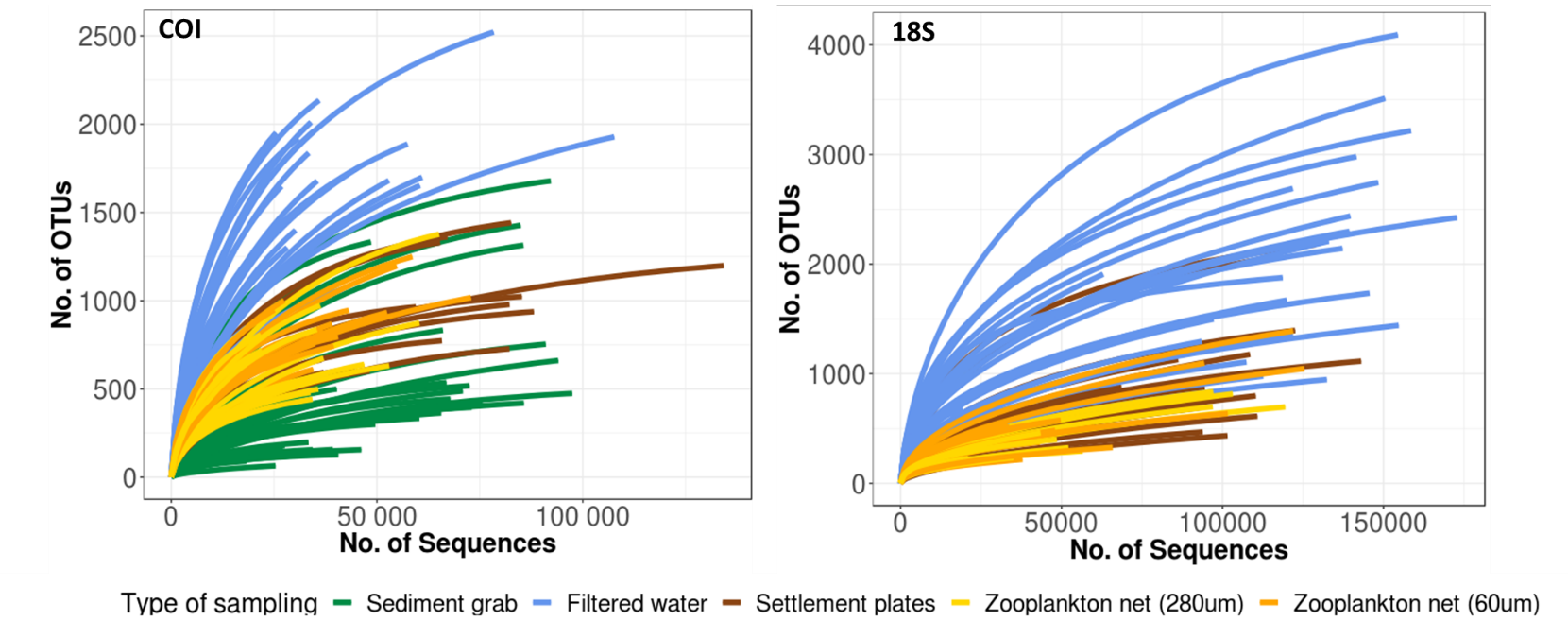

### Appendix 3

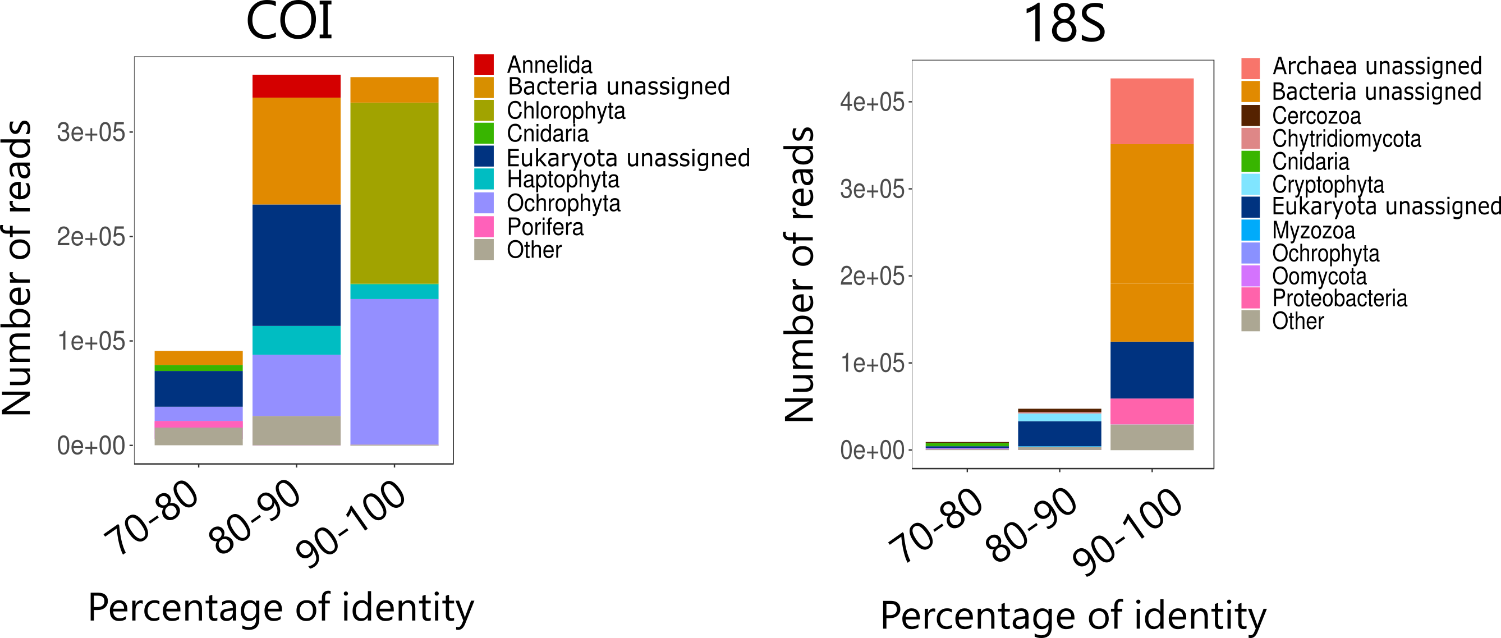

### Appendix 4

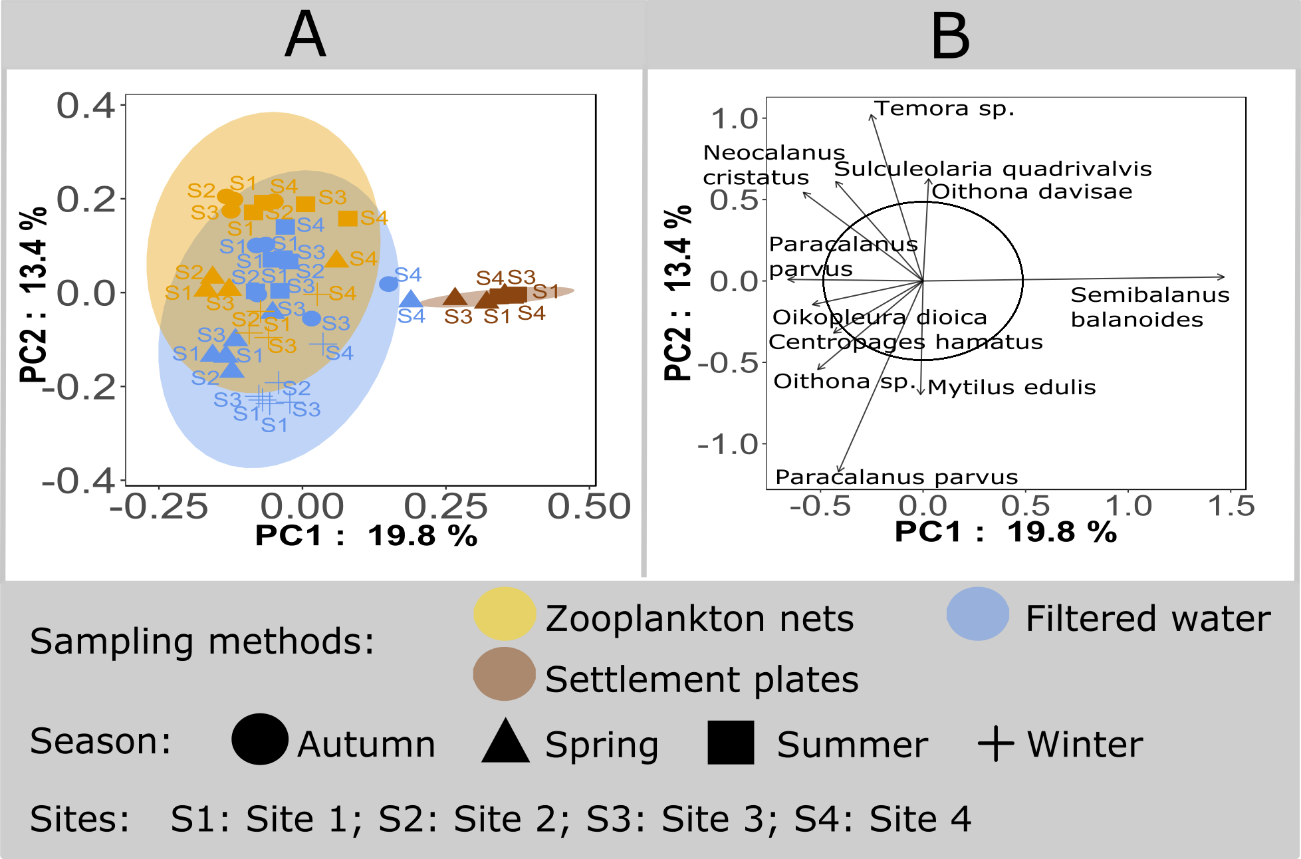

### Appendix 5

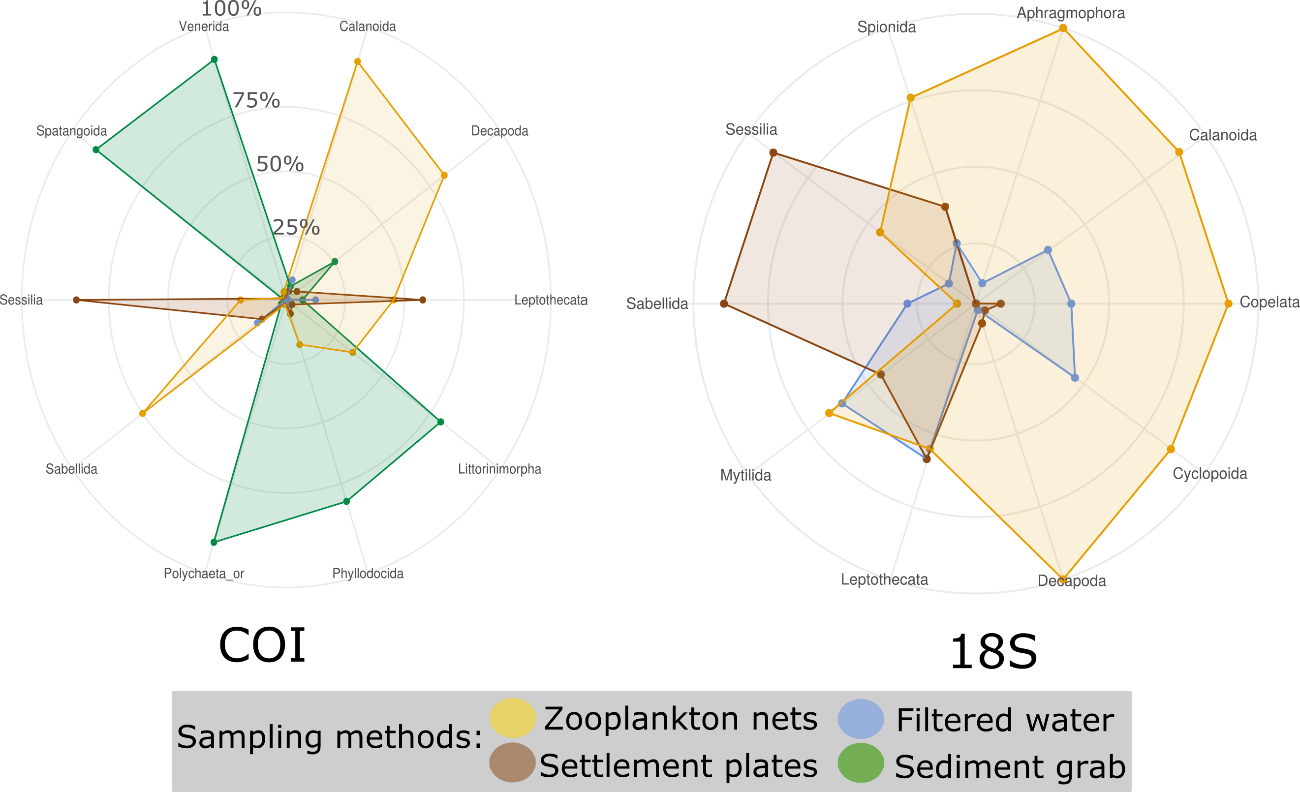

### Appendix 6

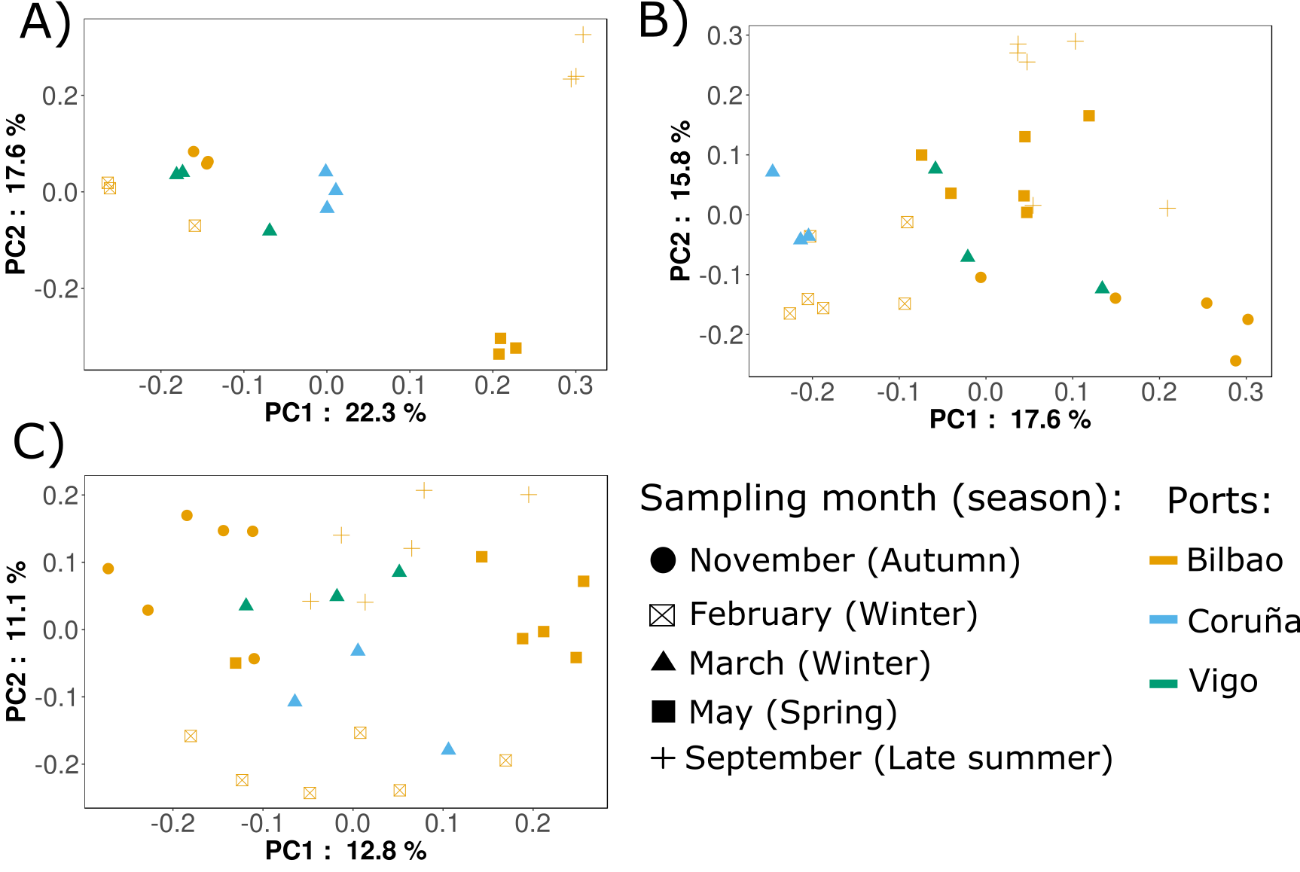
